# Functional In vivo Single-cell Transcriptome (FIST) Analysis Reveals Molecular Properties of Light-Sensitive Neurons in Mouse V1

**DOI:** 10.1101/382002

**Authors:** Jianwei Liu, Na Pan, Le Sun, Mengdi Wang, Junjing Zhang, Zhentao Zuo, Sheng He, Qian Wu, Xiaoqun Wang

## Abstract

Vision formation is classically based on projections from the retinal ganglion cells (RGC) to the lateral geniculate nucleus (LGN) and the primary visual cortex (V1). Although the cellular information of the retina and the LGN has been widely studied, the transcriptome profiles of single neurons with specific functions in V1 still remain unknown. Some neurons in mouse V1 are tuned to light stimulus. To determine the molecular properties of light-stimulated neurons in layer 2/3 of V1, we developed a method of functional *in vivo* single-cell transcriptome (FIST) analysis that integrates sensory evoked calcium imaging, whole-cell electrophysiological patch-clamp recordings, single-cell mRNA sequencing and three-dimensional morphological characterization in a live mouse, based on a two-photon microscope system. In our study, 58 individual cells from layer 2/3 of V1 were identified as either light-sensitive (LS) or non-light-sensitive (NS) by single-cell light-evoked calcium evaluation and action potential spiking. The contents of every single cell after individual functional tests were aspirated through the patch-clamp pipette for mRNA sequencing. Furthermore, the three-dimensional (3-D) morphological characterizations of the neurons were reconstructed in the live mouse after the whole-cell recordings. Our sequencing results indicated that V1 neurons with high expression of genes related to transmission regulation, such as *Rtn4r*, *Nr4a1,* and genes involved in membrane transport, such as Na^+^/K^+^ ATPase, NMDA-type glutamatergic receptor, preferentially respond to light stimulation. Our findings demonstrate the ability of FIST analysis to characterize the functional, morphological and transcriptomic properties of a single cell in alive animal, thereby providing precise neuronal information and predicting its network contribution in the brain.

## INTRODUCTION

Visual perception involves the activity of neurons in the cerebral cortex. The ‘retino-geniculo-cortical’ pathway indicates the best-known route for visual information ^1^. In the rodent primary visual cortex (V1), pyramidal cells together with inhibitory neurons confer highly specific visual features, such as light sensitivity and perceptual discrimination ^2,3^. It has been found that pyramidal cells in layer 2/3 with similar light orientation preferentially form synapses with each other ^4^. Specific types of interneurons in V1, such as parvalbumin (PV)- and somatostatin (SST)-positive interneurons, are also essential for modifying neuronal feature selectivity and improving perceptual discrimination ^5,6^. Recently, studies have revealed that direction-selective retinal ganglion cells (RGCs) deliver direction-tuned and orientation-tuned signals to superficial V1 ^7^. However, the molecular properties of the light-sensitive neurons in the V1 cortex still remain unclear. High-throughput single-cell RNA sequencing (scRNA-seq) has been applied to identify significant and divergent transcriptional responses of V1 neurons to sensory experience ^8^. However, since this approach was based on dissected tissue and isolated single cells, it is difficult to correlate the genetic profiles with the real-time stimulus-response and the electrophysiological characteristics of these cells.

The patch-seq technique, a method that combines whole-cell electrophysiological patch clamp recording and single-cell RNA sequencing, was presented to link the molecular profile to its corresponding electrophysiological and morphological counterparts in individual neurons ^9^. At present, the patch-seq technique has been widely used on mouse brain slices ^10^ or single neuron in culture ^11–13^. Since the physical environment during whole-cell recording on brain slices is different from the *in vivo* environment and the local circuitry related to acute stimulation is not able to be tested in the brain slices, a new technique to achieve patch-seq on a live animal during behavior tasks and electrophysiology recordings is required to be developed for detecting functional transcriptional responses in single cells.

We thus set out to develop a method for functional *in vivo* single-cell transcriptome (FIST) analysis for combining intracellular calcium imaging, *in vivo* whole-cell patch clamp recording, and high-quality RNA sequencing of individual neurons at layer 2/3 of the mouse V1 cortex, while the mouse was stimulated via light grating under light anesthetization. The FIST analysis identified the molecular biomarkers that were involved in the signal pathway of light sensitivity in V1. This technique could uncover the linkage from the cellular transcriptome profile to the acute sensory-evoked systematic response, and decode the neuronal physical execution at a gene transcription level. Thus, FIST technique is suit to discover the molecular elements that regulate the *in vivo* individual response, the cellular morphology and neuronal excitability.

## RESULTS

### Two-photon based in vivo calcium imaging of layer2/3 neurons in V1 of light stimulated mouse

In FIST system (**Fig. 1a**), a recording chamber was cranially fixed with a 1-mm^2^ window opened at the site of V1 (**Fig. 1b**). This method is largely applied to cells that are located near the surface of the cortex, at a depth of 100-200 μm (**Fig. 1c**). Guided by improved, two-photon microscopy, the new protocol is used to record intracellular calcium activity and whole-cell electrophysiological properties (**Fig. 1d-e**) and to further map the *in vivo* neuronal morphology (**Fig. 1f**). Additionally, this protocol can also be applied to record the intracellular calcium spikes in response to light stimulation followed by collection of the contents of the responding neuron for single-cell RNA sequencing (**Fig. 1g-i**).

**Figure 1:**
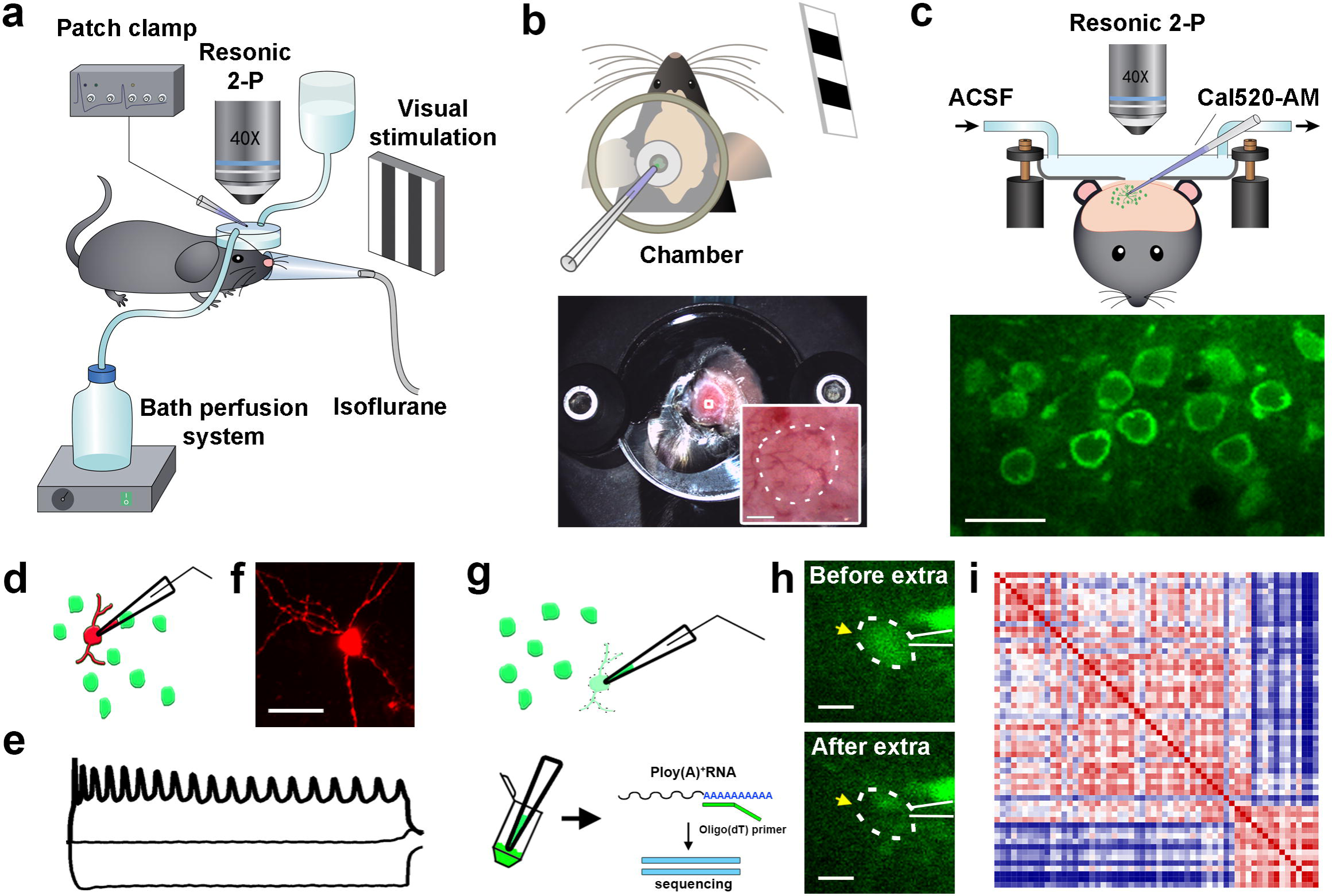
Schematics of the experimental approach of in vivo FIST analysis. **(a)** Overall view of the experimental arrangement of *in vivo* physiological recording with screening-evoked light stimulation. **(b)** Schematic of the recording chamber positioned at the V1 site of mouse. Bottom: top view of the chamber fixed at the V1 region. Insert: the cranial window (circled with a dotted line) opened at the center. Scale bar: 1 mm **(c)** Side view of the two-photon imaging setup for the *in vivo* patch-clamp recording and imaging. Bottom panel: Cal-520 AM labeled neurons. Scale bar, 20 μm. **(d)** Schematic of the *in vivo* whole-cell patch clamp. **(e)** Representative action potential firing recording of the *in vivo* patch clamp. **(f)** Imaging under the two-photon microscope. Red, staining for Texas Red. Scale bar, 20 μm. **(g)** Schematic of soma extraction of the target neuron for scRNA sequencing. **(h)** Example of a neuron before and after soma extraction. The dotted line indicates the location of the soma and the electrode is shown as white lines. Scale bar, 20 μm. **(i)** Schematic of cell transcriptome correlation by scRNA-seq analysis.

The cells at layer 2/3 of the V1 cortex were labeled with the Ca^2+^ indicator Cal-520 AM (**Fig. 2a and Supplementary Fig. 1**), which shows sufficient sensitivity for the detection of a single action potential and high signal-to-noise ratio ^14^. The sensory-induced calcium signal can be legibly detected in V1 cortical neurons of a lightly anesthetized mouse ^7^. The right eye of the mouse was given a 5-s visual stimulation of black-white drifting square-wave grating, with an inter-stimulation interval of 15 s and 5 repetitions of the visual stimulation in total. The calcium signals at a depth of ~120 μm in the V1 cortex were detected and recorded at a frequency of 30 frames/s, via the two-photon microscope equipped with two photomultiplier tubes (PMT) (**Fig. 2b**). The calcium response to the 5-time repeated visual stimulations was calculated immediately by manual matlab program (**Fig. 2c-d**). We defined a positive calcium response as having a calcium amplitude above the threshold (SNR = 2) at the moment of light stimulation. A cell showing ≥4 positive calcium responses in one round of testing with 5 repetitions of light stimulation was classified as a light-sensitive (LS) cell (**Fig. 2c-d, Cell 1 and Cell 2**). Non-light-sensitive (NS) cells were the ones with ≤ 1 positive response to the light stimulation. The identified cells were further recorded, imaged and soma extracted for RNA sequencing (**Supplementary Video 1**).

**Figure 2:**
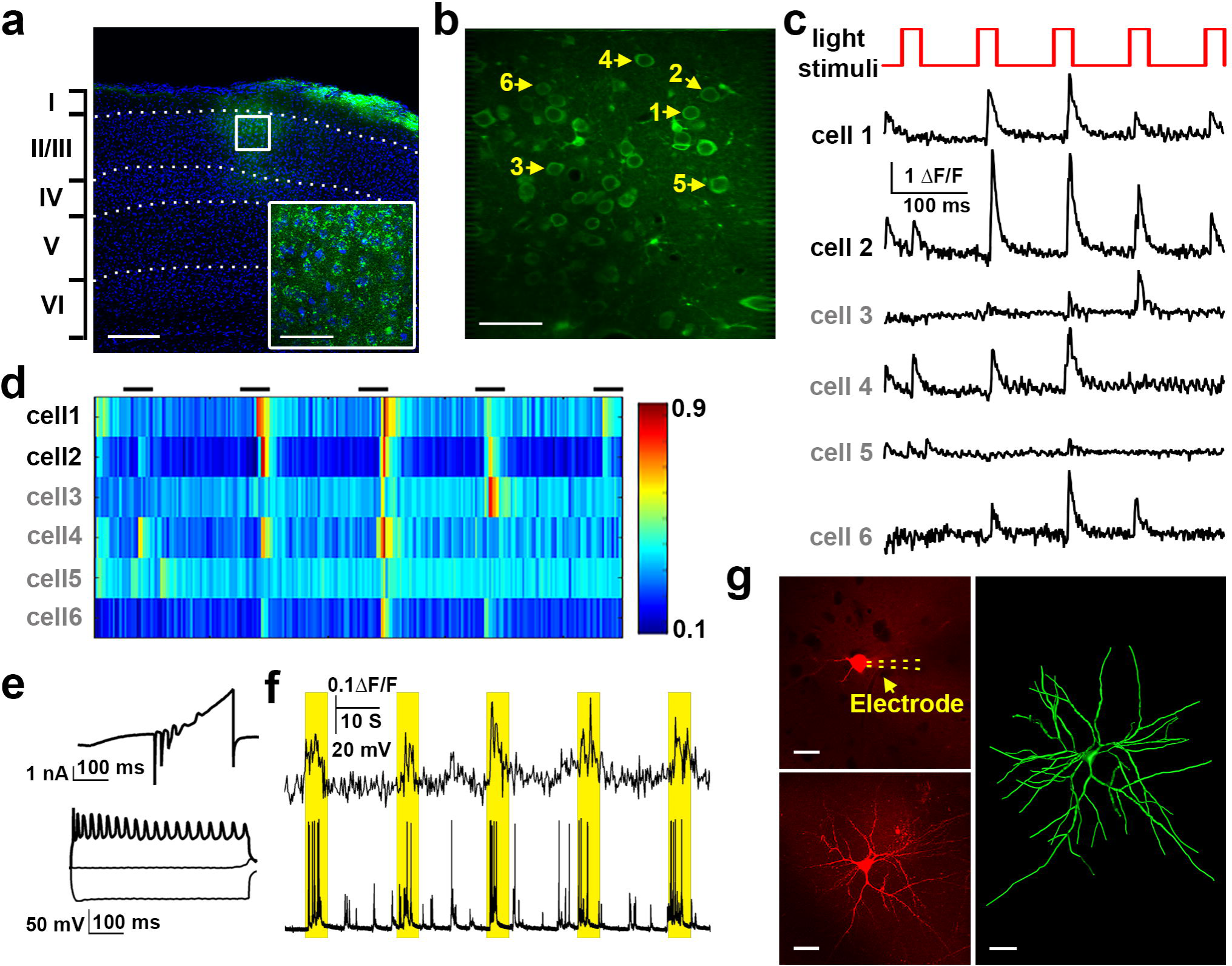
In vivo recording of light-sensitive neurons in layer 2/3 of V1. **(a)** Cal-520 AM labeled neurons in layer 2/3 of V1. Green, Cal-520 staining; blue, DAPI; white dotted line, laminar delimitation. Scale bar, 200 μm; scale bar of insert, 50 μm. **(b)** Calcium imaging of Cal-520 AM labeling, with yellow arrows indicating the six target neurons. Scale bar, 50 μm. **(c)** Calcium response to light stimuli of six neurons in (**b**). Top panel (red): visual stimulation sequence starts with a stationary period of square-wave gating for 5 s, with an inter-pulse interval of 15 s. Cells 1 and 2 (black label) were defined as light sensitive and cells 3-6 (gray label) were excluded according to our inclusion criteria described in the methods. **(d)** Heat map of the calcium responses to light stimulation of the six neurons in (**c**) (black bar). **(e)** Electrophysiological recording of a light-sensitive neuron. Upper panel: patterned action potential evoked by a stepped 500-ms current injection (−40 pA, 0 pA and 40 pA). Bottom panel: whole-cell current evoked by a 450 ms ramp voltage from −90 mV to +60 mV. **(f)** Representative dual recording of response (upper panel) and action potential firing (bottom panel) of one light-sensitive neuron. The yellow rectangle indicates the light stimulation period. Scale bar, 20 μm. **(g)** Morphology of a patch clamp recorded neuron using resonic two-photon imaging. Red, Texas Red perfused into the cytoplasm through the electrode (yellow dotted line); green, 3-D reconstruction of the neuron. Scale bar, 20 μm.

## In vivo electrophysiological and morphological analysis of LS neurons

Electrophysiological recording *in vivo* was applied to confirm the neuronal light sensitivity. We used the ‘red’ channel for Texas Red fluorescence (620/60 nm) to trace the electrode filled with Texas Red and confirm the located neuron by its shadow. As the patch-clamp pipette (with a resistance of 7-10 MΩ) was approaching a target cell, we continuously ejected the solution containing Texas Red and Oregon Green BAPTA-1 (OGB-1) from the tip with positive air pressure. When the electrode tip was sufficiently close to the target cell, we stopped ejection of the dye and applied negative air pressure to establish tight membrane sealing (> 1 GΩ), which was sustained for at least 2 minutes. A subsequent negative pressure was applied to rupture the membrane to form the whole-cell configuration, with an indication that the Texas Red diffused into the cytoplasm (**Supplementary Video 2**). A ramped I-V curve was applied to test the whole-cell current, including the inward sodium/calcium current and the outward potassium current (**Fig. 2e-upper panel**). The action potential firing was evoked by a stepped current injection (**Fig. 2e-bottom panel**). The calcium spikes and action potential were recorded at the same time as the light stimulation *in vivo*, showing the synchronic electrophysiological responses of a V1 cortical cell in response to light (**Fig. 2f and Supplementary Video 3**). This finding indicates that intracellular calcium imaging and analysis is a reliable method to identify neural activities *in vivo*.

Else, under the whole-cell patch clamp configuration, the Texas Red diffused into the cytoplasm (**Fig. 2g-left panels**). The morphology of the target neuron could immediately be imaged under the ‘red’ channel for Texas Red fluorescence in the two-photon microscope (**Supplementary Fig. 2**). The three-dimensional cellular morphology and structure of neurons were reconstructed using the z stack images and Imaris (**Fig. 2g-right panel and Supplementary Video 4**).

## Singe-cell RNA sequencing of somatic aspirates

For single-cell RNA sequencing, the autoclaved RNase-free pipette (with a resistance of 2-3 MΩ) was made to approach a target cell after calcium spike recording as described above. When the electrode tip was sufficiently close to the target cell, we quickly sucked the soma into the pipette and collected the sample immediately after confirming that the soma entered by breaking the pipette tip (**Fig. 1h-i and Supplementary Video 1**). The internal pipette solution containing the lysis buffer had an optimal volume of less than 1 μL ^9^.

After harvesting the cell contents, the single-cell mRNA was converted to cDNA and used to generate sequencing libraries following the protocol of Smart-seq2. Cells with a low cDNA concentration (< 1,000 pg/μL) (**Supplementary Fig. 3a**) and poor sequencing quality (< 1,000 genes) (**Supplementary Fig. 3b**) were excluded from further analysis (3/63 cells). Cells with a high mitochondrial RNA fraction (> 25%) were also omitted (2/60 cells) (**Supplementary Fig. 3c**). In the end, 58 out of 63 cells (92.1%) from 24 mice passed the quality control according to our inclusion criteria, including 44 light-sensitive (LS) cells and 14 non-light-sensitive (NS) cells. We detected approximately 5,700 genes for which the fragments per kilobase of transcript per million total reads (FPKM) was higher than 1, and 4,500 genes for which the FPKM was higher than 5 per cell (**Supplementary Fig. 3d**), with a median Pearson correlation of 0.59 for LS cells and 0.49 for NS cells (**Supplementary Fig. 3e**).

**Figure 3:**
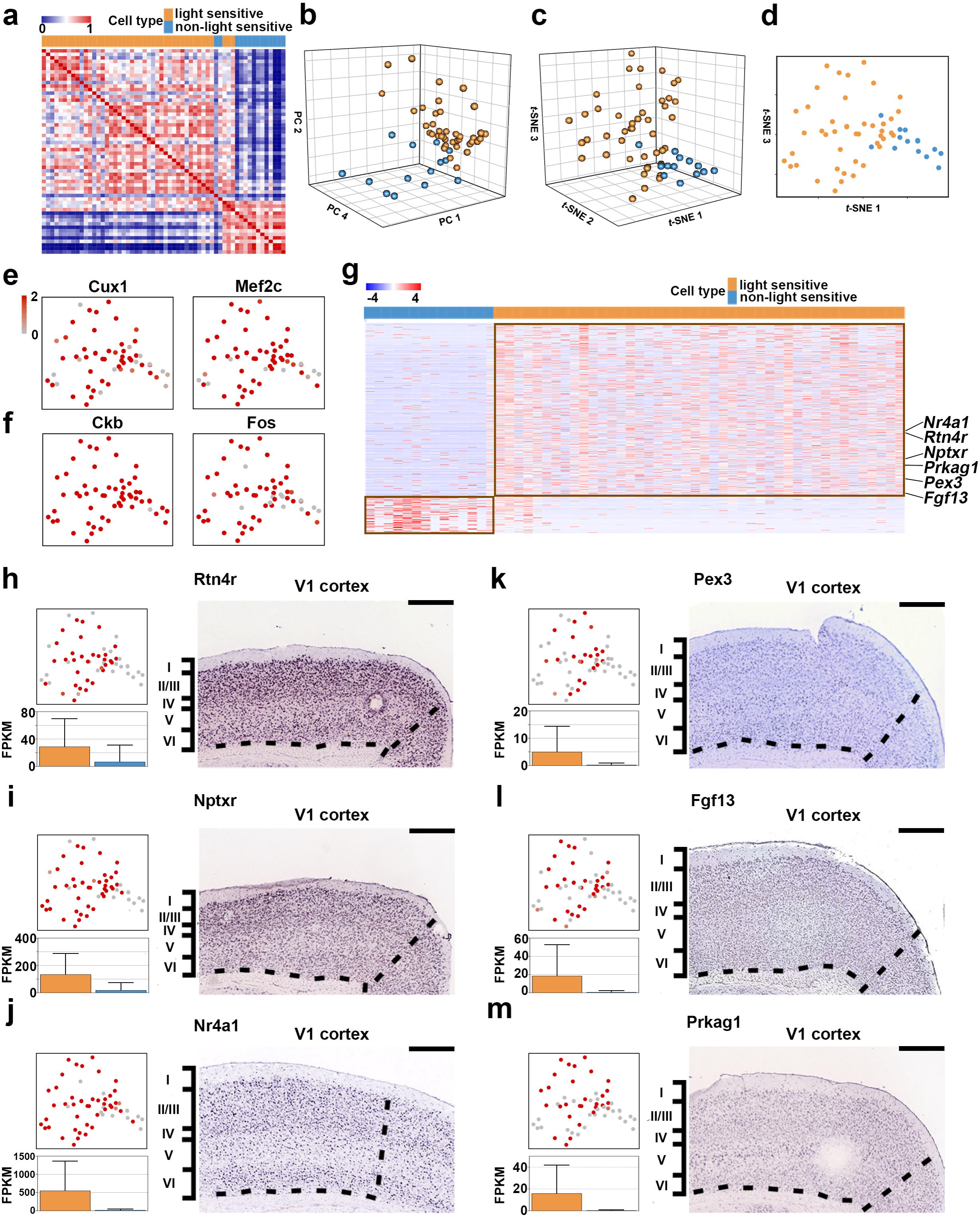
Single-neuron transcriptome profile. ** (a)** Pearson correlation heatmap of 58 cells from V1 of 24 mice. Clustering analysis separates light-sensitive (LS) and non-light-sensitive (NS) neurons. ** (b)** Three-dimensional principal component analysis (PCA) of expression profiles of 20183 genes detected in 58 sampled cells. (**c-d**) Three-dimensional representation of 58 samples using t-SNE (**c**), with two-dimensional t-SNE drawn by the first and third components in (**d**). Cells are colored according to clustering as LS and NS cells. **(e-f)** Marker gene expression by cortical layer 2/3 neurons (**e**) and in primary visual cortical cells (**f**) overlaid onto the two-dimensional t-SNE. **(g)** Heatmap of log2 transformed gene expression of differentially expressed genes. Genes with P < 0.05 and log2foldchange > 1 or log2foldchange < −1 were identified as differentially expressed genes. The differentially expressed genes were ranked in descending order based on their log2foldchange value. **(h-m)** Identification of genes expressed differentially between LS and NS neurons. The gene expression was overlaid onto the two-dimensional t-SNE, colored according to the FPKM normalized gene expression level. Expression quantitation is represented on the histogram as mean ± s.d. The expression pattern in V1 is shown as an ISH image (from Allen Brain Atlas). Scale bar, 420 μm.

### Mapping neuronal identities on FIST data sets

Out of 58 screened cells, two groups of cells were demonstrated by Pearson-correlation-based classification, which were mostly consistent with our functional definition of the LS and NS cells (**Fig. 3a**), suggesting that cells with the same functions may have similar transcriptomic profiles. After dimensionality reduction of the expression profile of 20,183 genes detected in 58 cells, LS- and NS-cells were clustered into two groups, using either principal component analyses (PCA) (**Fig. 3b**) or t-distributed stochastic neighborhood embedding (t-SNE) (**Fig. 3c-d**). Transcriptome analysis indicated that all the cells are neurons because they expressed marker genes of pyramidal cells (*Neurod1*, *Neurod2*, and *Emx1*) or interneurons (*Calb1*, *Cck*, and *Slc6a1*) (**Supplementary Fig. 4a**). These neurons demonstrated a high level of expression of the genes *Cux1* (Cut like homeobox 1), *Cux2* (Cut like homeobox 2), and *Mef2c* (Myocyte-specific enhancer-binding factor 2) (**Fig. 3e and Supplementary Fig. 4b**), indicating that these are cells located in cortical layer 2/3 of adult mice ^15–17^. Moreover, we also observed high expression in these cells of *Ckb* (Creatine kinase B), *Fos* (c-Fos) and *Junb* (Jun-B) (**Fig. 3f and Supplementary Fig. 4c**), which were reported in mouse V1 in a recent study using high-throughput scRNA-seq ^8^. Together, the expression of marker genes in all cells suggests that FIST analysis provides high-quality transcriptome data and that the samples we have collected are layer 2/3 V1 neurons.

**Figure 4:**
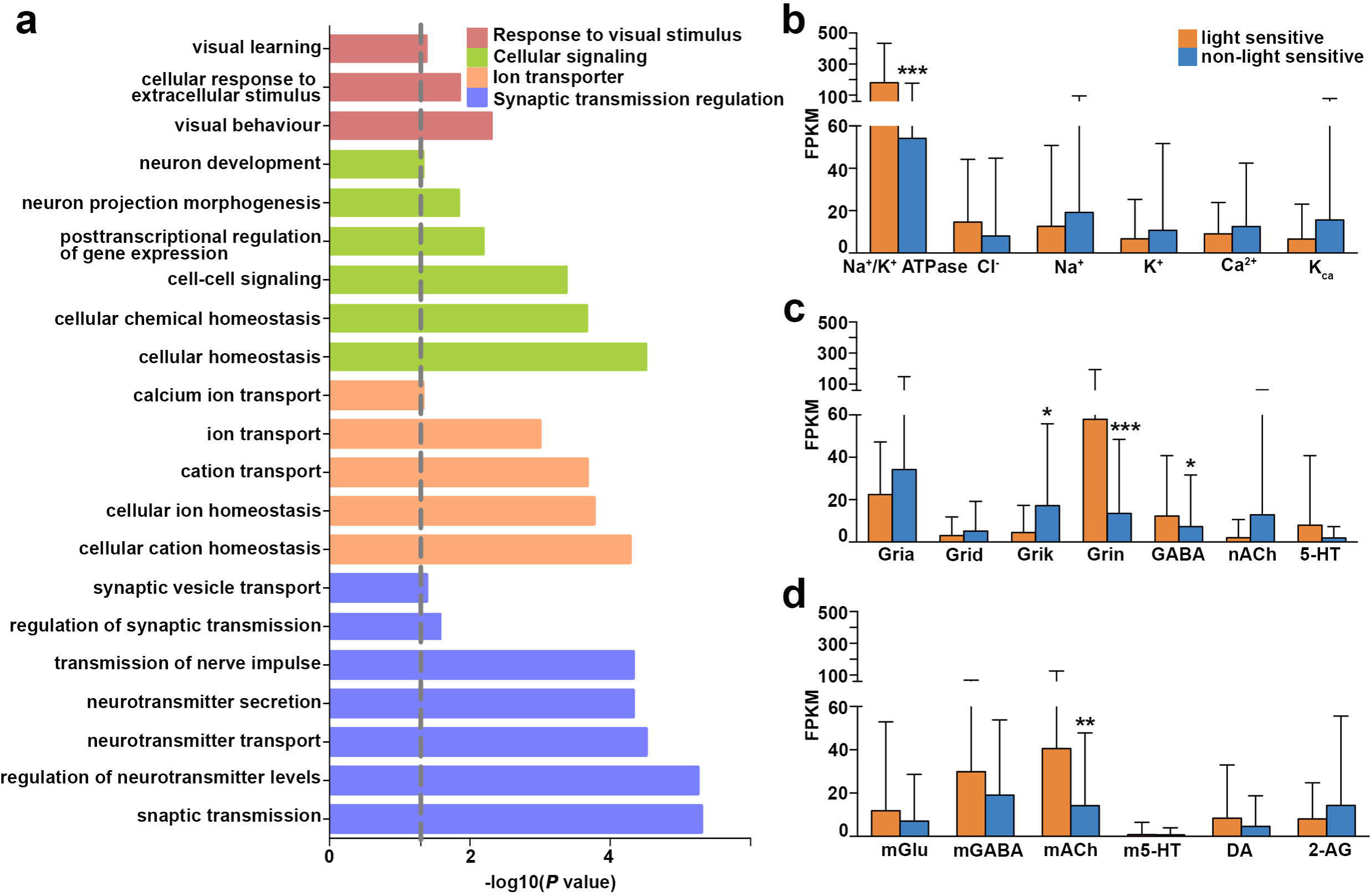
Enriched gene ontology analysis. (**a**) Histogram of the enriched GO terms of differentially changed genes from the pairwise comparison of light-sensitive and non-light-sensitive cells. Dotted line, p = 0.05. (**b-d**) Quantitative expression of ion channel and membrane receptor genes in light-sensitive and non-light-sensitive cells. Genes include Na^+^/K^+^-ATPase, voltage-gated Cl^-^, Na^+^, K^+^, Ca^2+^, and Ca^2+^-activated K^+^ ion channels in (**b**); ligand-gated ionotropic channels of AMPA-, delta-, kainite- and NMDA-type glutamatergic receptors (Gria, Grid, Grik, and Grin, respectively); GABAergic (GABA), nicotinic acetylcholinergic (nACh) and serotonergic (5-HT) receptors in (**c**); metabolic glutamatergic (mGlu), GABAergic (mGABA) receptors, 5-HT, muscarinic acetylcholine (mACh), dopamine (DA) and 2-arachidonoylglycerol (2-AG) receptors in (**d**). Error bar, mean ± s.d. The significance was estimated based on the corrected P value using the unpaired t-test, * p < 0.05, ** p < 0.01, *** p < 0.001.

To identify the molecular properties involved in the light sensitivity of neurons, the differentially expressed genes (DEGs) between the LS neurons and NS neurons were found (**Fig. 3g**). Genes with P < 0.05 and log2FoldChange > 1 or log2FoldChange < −1 were identified as DEGs. In total, we screened 550 DEGs, out of which 450 genes were more highly expressed in LS neurons and 100 genes were more highly expressed in NS neurons. In our data, the LS neurons expressed much higher levels of *Rtn4r* (also called *Ngr1*, the nogo receptor 1)*, Nptxr* (the neuronal pentraxin receptor) and *Nr4a1* mRNA than did NS neurons (**Fig. 3h-j**). Among the up-regulated genes, *Rtn4r* was reported to mediate intralaminar synaptic connectivity of visual cortical interneurons or a subclass of PV^+^ interneurons ^18,19^, which markedly sharpened grating orientation tuning and enhanced direction selectivity of nearby neurons ^5^. Additionally, *Nptxr* has been hypothesized to be involved in activity-dependent synaptic plasticity ^20,21^, as well as synaptic maturation through PV^+^ interneurons in V1 ^22^. The Nptx1/2-expressing RGCs project to the dorsal lateral geniculate nucleus (LGN) neurons ^23^ and precisely map the visual cortex pattern ^24^. Thus, the high expression of *Nptxr* would indicate that it plays a key role in the visual pathway of light experience. Additionally, genes that influence transmission strength and synapse plasticity during the individual selection of superior neurons and elimination of inferior neurons also showed differential expression between the LS- and NS-groups, such as *Nr4a1* ^25,26^. In summary, our data pool showed that some genes of known function are highly expressed in V1 neurons in response to light stimulation, indicating that synapses of light-active neurons in mouse V1 are possibly strengthened and maintained through competitive processes that require specific transcriptional regulation.

In addition to well-studied genes involved in the visual system, we also found that *Pex3* (peroxisomal biogenesis factor 3), *Fgf13* (fibroblast growth factor 13) and *Prkag1* (protein kinase AMP-activated non-catalytic subunit gamma 1) have potential roles in regulating the light sensitivity of neurons in V1 (**Fig. 3k-m**). The genes are either contributing to the transmitter vesicle assembling ^27^ or responsible for axonal terminal growth ^28^, or they may even be involved in manipulating the AMPK signal pathway ^29^, suggesting that LS neurons might be more compatible for light-evoked input transduction and signal processing in V1.

### Molecular candidates to determine light sensitivity of neurons in V1

To further investigate the specific biological properties of LS neurons, Gene Ontology (GO) analysis was applied to the 200 most highly expressed genes of these cells. Genes were enriched into 18 GO terms and divided into 4 clusters, including response to the visual stimulus, cellular signaling, ion transporters and synaptic transmission regulation (**Fig. 4a**). Neurons dynamically tune their excitability via the integration of transmembrane ion channels and transporters and accomplish afferent and efferent communications by receptor-dependent transmitter release ^30–32^. Most of the information regarding transcriptome profiles of transmembrane ion channels, ligand-gated inotropic channels and metabotropic (G protein-coupled) receptors of neurons were collected in our FIST data pool (**Fig. 4b-d**). We used the unpaired Welch’s two-sample t-test to determine if two groups were significantly different from each other. Among the transmembrane proteins, the components that generate an action potential, including potassium, sodium and calcium channels, are consistent in the LS and NS groups. However, the expression levels of the Na^+^/K^+^-ATPase, ionotropic glutamate and GABA receptors and metabolic acetylcholine receptors were higher in LS cells than in NS cells.

The Na^+^/K^+^-ATPase, which also functions as a signal transducer to regulate intracellular calcium in the retina ^33^ and visual cortex ^34^, showed higher mRNA expression in LS neurons than in NS neurons (P < 0.00001) (**Fig. 4b**). Considering that the Na^+^/K^+^-ATPase participates in intracellular calcium signaling ^35^, our data suggest that light stimulation triggers more calcium oscillations in LS neurons than in NS neurons. In addition, the subunits of the NMDA receptor (P < 0.00001) and kainite receptor (P < 0.05) are differentially expressed in the LS and NS groups. On the other hand, the majority of ionotropic GABA receptors also have different expression levels in LS and NS neurons (P < 0.05) (**Fig. 4c**). It is well known that glutamatergic transmission mainly relies on the N-methyl-D-aspartate (NMDA) receptor, α-amino-3-hydroxy-5-methyl-4-isoxazolepropionic acid (AMPA) receptors, delta receptors and kainite receptors expressed in neurons ^36^. Previous and current studies have mentioned the mechanisms of NMDA receptor-induced LTD and LTP that account for key aspects of experience-dependent synaptic modification in the visual cortex ^37,38^, as well as the GABAa receptor-dependent intracortical inhibitory inputs, are suggested to be involved in regulating the orientation tuning of the spike responses in the visual cortex ^39,40^. These results suggest that gene-transcription-regulated synaptic transmission dynamically promotes the light sensitivity of a subtype of V1 neurons. We also observed that the NS neurons lack appreciable mRNA expression of the muscarinic acetylcholine receptors (mAChR), which have a significant expression in LS neurons (P < 0.01) (**Fig. 4d**). Preliminary studies have reported that ACh application could increase neuronal orientation and direction selectivity in cat V1 ^41,42^; moreover, recent studies have also mentioned that the mAChR could regulate the neuronal activity in V1 to improve light contrast sensitivity in monkeys ^43^, which was consistent with our sequencing data.

## DISCUSSION

*In vivo* FIST analysis is an approach that enables functional cellular activity recording and high-quality RNA-seq of individual neurons in live animals. It generates a comprehensive profile of an individual neuron, including cellular physiology (calcium imaging and electrophysiological activities), morphology and transcriptional responses at the single-cell level.

Using this technique, we have illustrated that light-sensitive V1 neurons are functionally and transcriptionally correlated, and we also identified several molecular markers of these cells. Similar with the high-throughout RNA sequencing data sets in mouse V1 after chronic light exposure ^8^, our transcriptome data showed that the sensory-stimulation-induced expression across most neurons contained regulator genes, such as, *Nr4a1* and *Fosb*. Else, the quickest transcriptional response known in any setting is the induction of immediate-early genes, including *Fos*, *Junb*, *Egr4* ^8^. In our data, these neuronal activity response genes showed significant high expression in LS than NS neurons. Furthermore, our transcriptome data also suggest that typical synaptic receptors might be involved in light-stimuli responses in V1 neurons. Additionally, the genes highly expressed by V1 neurons involved in the light response (such as *Nptxr*) functionally related to the genes expressed in the retina and LGN (such as *Nptx*), indicating the transcriptomic feature along the ‘retino-geniculo-cortical’ pathway.

There are currently two published protocols for Patch-seq by Cadwell et al. and Fuzik et al. ^9,10^. However, they have not connected the single cell transcriptome profiles with the sensory-evoked function *in vivo*. Our whole-cell recording in LS neurons indicated that the action potential firing, followed by the intracellular calcium evaluation, was strongly associated to the light stimulation on the eye. Our FIST analysis data suggested the rapid physiological response of LS neurons might attribute to the higher expression level of transmembrane protein, such as Na^+^/K^+^-ATPase, ionotropic glutamate and GABA subtype receptors. It is suggested the neurons that possessed stronger transmembrane conduction, including the ion transporting, synaptic transmission, and intracellular calcium variation, would prefer to be selected as the sensory-evoked neurons, like the LS neurons in our study.

In summary, our approach can be used broadly to combine the information of *in vivo* context-evoked physiological records, cell morphologies and transcriptional responses of single neurons in live animals, which will be applied in behavioral process and disease models of animals, including rodents and primates. Performing unbiased, whole-genome transcriptome analysis and physiological characterization of individual neurons of living animals with FIST analysis might help to resolve long-standing questions and open new directions of investigation in neuroscience.

## METHODS

### Surgical procedures for in vivo experiments

Experiments were performed on adult male C57/B16 mice (age 9-11 weeks). Animals were maintained on a 12 h light/12 h dark cycle. Mice were anesthetized using an isoflurane-oxygen mixture (3.0% (v/v) for induction and 1.5% (v/v) for maintenance) and kept in a prone position in a stereotaxic apparatus with a heating pad sett to 37-38 °C. After a midline scalp incision, a custom-made plastic chamber was then fixed to the skull with instant adhesive (ALTECO^®^) and dental cement over the left primary visual cortex (V1) according to stereotaxic coordinates (center: ∼2.7 mm lateral and 3.5 mm posterior to the bregma). A small, square craniotomy (2 mm × 2 mm) was made with a pneumatic dental drill. The dura mater was carefully removed by with forceps. Afterwards, the mouse was transferred to and fixed on the recording setup, with light anesthetization with an isoflurane-oxygen mixture of 0.5-1% (v/v). The recording chamber was perfused with normal artificial cerebral spinal fluid (ACSF) containing 126 mM NaCl, 3 mM KCl, 1.2 mM NaH_2_PO_4_, 2.4 mM CaCl_2_, 1.3 mM MgCl_2_, 26 mM NaHCO_3_, and 10 mM D-glucose (pH 7.4 when bubbled with 95% oxygen and 5% CO_2_). The temperature of the mouse was kept at ~37 °C throughout the experiment.

### Dye loading

The highly sensitive fluorescent Ca^2+^ indicator Cal-520 AM (Santa Cruz Cat. no. sc-477280) was used for multicellular bolus loading in the V1. Cal-520 AM was dissolved in DMSO with 20% Pluronic F-127 to a final concentration of 250 μM with loading buffer (150 mM NaCl, 2.5 mM KCl, 10 mM HEPES, PH = 7.4) for bolus loading. A micropipette was filled with this solution and inserted coaxially into the cortex and down, reaching a depth of 120 μm below the cortical surface (**Fig. 1c**). A pressure pulse [15 min, 0.2 bar (1 bar 100 KPa)] was applied to the pipette to eject the dye-containing solution ^44^. The dye of Texas Red 3000 was also dissolved (0.1% solution) in the standard pipette solution and applied through micropipettes similar to those used for injections of AM indicator dyes. We performed the Ca^2+^ imaging ~1 hour after dye injection, and the imaging lasted for up to 7 hours.

### Visual stimulation

Visual stimuli were presented as described previously ^2^. Briefly, stimuli were displayed with correction on an LCD monitor (7 inch, 1024×768, 60 Hz refresh rate) placed approximately parallel to and 16 cm from the right eye of the mouse, with the visual angle subtended by the monitor (± 23.5º azimuth and ± 26.9º elevation). A paperboard trapezoid cylinder placed between the eye and the screen was used to prevent stray light. The stimuli covered the full screen with black-white drifting square-wave grating presented on a dark gray background. The visual stimuli were generated by a program written in C#. Each trial of visual stimulation sequence started with a stationary six periods of square-wave grating for 1 s. The luminances of black, white and gray were 3.64, 0.11 and 0.12 cd/m^2^, respectively. The grating drifted for 5 s (spatial frequency 0.06 cycles per degree, drifting speed 1.2 cycles per second), with an inter-stimulus interval of 15 s. Each stimulus was repeated 5 times. Neurons that were responsive to the grating less than 4 times were excluded from our data.

### In vivo electrophysiology and RNA extraction

To collect an initial dataset of morphology and electrophysiology of neurons, the patch pipette solution contained the following (in mM): 130 K-gluconate, 16 KCl, 0.2 ethylene glycol-bis-(b-aminoethylether)-N,N,N,N-tetraacetic acid (EGTA), 2 MgCl_2_, 10 HEPES, 4 Na_2_-ATP and 0.4 Na_6_-GTP. Approximately 0.1% Texas Red 3000 as a tip indicator was added to the electrode and diffused within the recording neuron during the whole-cell recording for further confirmation staining. Patch-clamp pipettes were pulled from borosilicate glass (1.5 mm OD × 0.84 mm ID, VitalSense Scientific Instruments) with electrode resistances ranging from 7 to 10 MΩ. Tight seals (> 1 GΩ) were obtained on cell bodies before rupturing the membrane with negative pressure. Whole cell recordings were conducted with the Axon 700B patch-clamp. The currents were typically digitized at 100 KHz, and macroscopic records were filtered at 2 KHz.

To obtain transcriptome data from single neurons, the internal pipette solution was modified as followed (in mM): 123 K-gluconate, 12 KCl, 10 HEPES, 0.2 EGTA, 4 MgATP, 0.3 NaGTP, 10 sodium phosphocreatine, 20 μg/mL glycogen, and 1 U/μL recombinant RNase inhibitor ^45^. Approximately 0.1% Texas Red as a tip indicator, and 6 μM calcium indicator dye Oregon Green BAPTA-1 (OGB-1) was added for membrane rupture and inner extraction. All neuron-related reagents were RNase-free. The electrode resistance ranged from 2 to 4 MΩ. The cell membrane was ruptured via negative pressure with a calcium influx indicated by OGB-1. The extracted soma of the neuron was confirmed as the electrode was pulled out of the tissue and then transferred to ice-cold lysis buffer immediately after breaking the tip within the tube. The sample was stored at −80 ºC for at most one week before the next step of processing.

### Two-photon imaging

Fluorescence was imaged with a two-photon microscope (Scientific Inc.) equipped with a mode-locked Ti:sapphire laser (MaiTai, Spectra-Physics) and a water immersion objective lens (Apo40×W/NIR, NA 0.8; Nikon). The emitted photons are split into two channels and detected by photomultiplier tubes (PMT), i.e., into a ‘green’ channel for Cal-520 AM and OGB-1 fluorescence (525/50 nm) and a ‘red’ channel for Texas Red fluorescence (620/60 nm). With this configuration of our system, one pixel corresponded to 0.31×0.31 μm^2^ in x,y-coordinates. A square region of 160 × 160 μm^2^ (512 × 512 pixels) was scanned at the maximum. The frame rate was 30 frames/s.

### Statistical analysis

The calcium response data were analyzed via ImageJ software (version 1.49g). The cell body pixels were selected manually for each target neuron, as well as the pixels surrounding the cell body as the background. Next, the ratio (∆F/F) of the stimulus-evoked change (∆F) to the average level (F) was calculated as the difference between the average fluorescence signals of the cell body and the background along with the time lapse; the ratio was then normalized with the mean fluorescence across the pixels of each neuron. The calcium spike was defined as an event when the ratio was higher than twice the non-spike level (SNR ≥ 2). The response magnitudes of light-evoked calcium spikes were defined as the mean value of the difference between the peak value during the post-stimulus period (5 s) and the basal average value during the pre-stimulus period (2-4 s), after 5 repetitions.

Electrophysiological data were analyzed with ClampFit software (version 10.0; Axon Instruments) and GraphPad Prism (v.6.0; GraphPad Software). A whole-cell current-voltage curve under a 450-ms ramp command from −90 mV to +60 mV was intended to test the whole-cell current of the neuron. The patterned firing spikes were evoked by a series of 500-ms current pulse injections from −40 pA to 40 pA (40 pA/increment). The field potential was recorded at a current clamp mode with no current injection.

Three-dimensional morphological reconstruction was conducted with Imaris software (version 7.6.0, Bitplane).

### Library construction and sequencing

We converted the RNA collected from the *in vivo* patch-clamped neurons to cDNA for generating the sequencing libraries following the protocol of Smart-seq2 ^46^. The primers for reverse transcription were referenced from Cadwell’s procedure ^9^. After 24 cycles of amplification, approximately 60 ng of purified cDNA was used to construct sequencing libraries using the commercial KAPA HyperPlus Kit protocol (KK8514). The PCR products with different index sequences were pooled together for purification and library construction. Quality control was performed on both the amplified cDNA and the final library using a Fragment Analyser (Thermo Fisher). The DNA was sequenced using an XTEN platform. Investigators were blinded to cell type during library construction and sequencing.

### Read processing and quantification of gene expression

Adapters and low-quality reads were trimmed using Python script AfterQC ^47^. Paired-end reads were aligned to the reference genome GRCm38 primary assembly ^48^ downloaded from ensemble using STAR (STAR 2.5.3a) ^49^ with default settings except for the use of setting output type (--outSAMtype) to sort SAM outputs by coordinates. Reads were then counted using featureCounts (featureCounts 1.5.3) ^50^. The R package, scater (Single-cell analysis toolkit for gene expression data in R, version 1.6.3) ^51^, was employed for quality control, normalization and data visualization. Gene expression was normalized to the number of fragments per kilobase million (FPKM value). Genes that did not have an expression value of at least 1 FPKM across all samples were excluded from further analysis. Moreover, only protein coding genes were considered for subsequent analysis. The filtered dataset contained 20,183 genes in total.

### Quality control

Two cells with cDNA concentration lower than 1,000 pg/μl were discarded. Another two cells with a mitochondrial DNA proportion greater than 25% were omitted. One cell with fewer than 1,000 expressed protein coding genes (which had an expression level of at least 1 FPKM) was excluded as well. The dimensionality of the dataset was then reduced from 63 cells to 58 cells.

### Dimensionality reduction techniques

For dimensionality reduction, principle component analysis (PCA) and t-distributed Stochastic Neighbor Embedding (t-SNE) were performed using function plotPCA from R package scater and function Rtsne (version 0.13), respectively. The dataset was projected onto 2- and 3-dimensional space with both methods.

### Heatmap of Pearson correlation matrix

Pearson correlation between light-sensitive and non-light-sensitive subsets was computed. A heatmap was generated using R package pheatmap (version 1.0.8).

### Differential gene expression analysis

Differential gene expression analysis was performed using R package Deseq2 (version 1.18.1) ^52^. Genes as P < 0.05 and log2FoldChange > 1 or log2FoldChange < −1 (higher absolute value means higher fold change of corresponding clusters) were identified as differentially expressed genes (DEGs). DEGs were ranked in descending order based on their log2FoldChange value. A heatmap of gene expression from 550 DEGs was generated using R package pheatmap (the first 450 and the last 100 DEGs).

### Identification of highly variable genes

The top 200 most expressed genes from the light-sensitive subset were identified using plotQC function from R package scater by setting type = “highest-expression”. Enriched Gene Ontology (GO) analysis of those genes was performed using the online tool, DAVID 6.7 ^53,54^. Function geom_bar from R package ggplot2 (version 2.2.1) was employed for drawing barplots.

### Ion channel and receptor-related gene analysis

Expression levels of the following ion channel and membrane receptor-related genes across all 58 samples were computed: Na^+^/K^+^-ATPase, voltage-gated Cl^−^, Na^+^, K^+^, Ca^2+^ and Ca^2+^-activated K^+^ (K_Ca_) channel subtypes; ligand-gated ionotropic channels of AMPA-, delta-, kainite- and NMDA-type glutamatergic receptors (Gria, Grid, Grik, and Grin, respectively); GABAergic (GABA), nicotinic acetylcholinergic (nACh) and serotonergic (5-HT) receptors; metabolic Glu (mGlu), GABA (mGABA), 5-HT, muscarinic acetylcholine (mACh), dopamine (DA) and 2-arachidonoylglycerol (2-AG) receptors. Barplots of each gene with the corresponding quantitative expression levels across all samples were generated using R function geom_bar. To determine if the two subsets were significantly different from each other, the t-test was employed. Because variances in the two datasets were unequal, the unpaired welch two-sample t-test was performed using R function t.test by setting var.equal = FALSE.

## ACKNOWLEDGMENTS

We thank Dr. Xiaowei Chen (Third Military Medical University, Chongqing, China) for discussion. We thank Dr. Lu Yang’s help on solving single-cell sequencing technical problems. This work was supported by National Basic Research Program of China (2017YFA0103303, 2017YFA0102601), the Strategic Priority Research Program of the Chinese Academy of Sciences (XDA16020601), the National Natural Science Foundation of China (NSFC) (91732301, 31771140) to X.W., (31600828) to N.P., (31671072) to Q.W., Youth Innovation Promotion Association CAS to Q.W.

## AUTHOR CONTRIBUTIONS

Q. W., S. H., and X.W. conceived the project, designed the experiments and wrote the manuscript. J. L. and N. P. conducted the animal surgery, calcium imaging and electrophysiology experiments. M. W. and Z. Z. performed single-cell RNA-seq and data analysis. J. L. and L. S. performed 3D reconstruction of neurons. J. Z. maintained the animals. All authors edited and proofed the manuscript.

## DATA AVAILABILITY

The scRNA-seq data are available in Gene Expression Omnibus under accession number GSE115997.

## Supplementary Figure 1: Confirmation of Cal-520 AM labeling in layer 2/3 of V1

Sagittal slice containing the V1 area. Left, Cal-520 labeled neurons in V1. Green, Cal-520 staining; blue, DAPI; white dotted line, V1 delimitation. Right, corresponding mouse brain map for V1.

## Supplementary Figure 2: 3-D animation of morphological reconstruction

Morphology of recorded neurons under the two-photon microscope. Red, Texas Red, scale bar: 20 µm; green, 3-D reconstruction of respective neurons; scale bar, 20 µm.

## Supplementary Figure 3: Quality control of cDNA libraries

1. Concentration of all collected samples. Only cells with more than 1,000 pg/μL cDNA were sequenced (cutoff denoted by the dashed line). The excluded samples are indicated in red.
2. Mitochondrial proportion of the samples screened in (**a**). The cells with higher than 25% mitochondrial RNA (red dots) were excluded from the sample pool (cutoff denoted by the dashed line).
3. Average library sizes generated from the samples screened in (**b**). Dashed line indicates threshold criteria used for sequencing. Poor quality libraries with lower than 1,000 genes (FPKM > 1) were excluded (red dots).
4. Number of genes detected per neuron using two different expression thresholds.
5. Pairwise Pearson correlation across all detected genes for light- and non-light-sensitive neurons.

## Supplementary Figure 4: Marker gene expression overlaid onto t-SNE map

1. Marker gene expression of cortical neurons including pyramidal cells and interneurons using two-dimensional t-SNE representation.
2. Cortical layer 2/3 marker gene expression overlaid onto the t-SNE map.
3. Reported enriched V1 gene expression overlaid onto the t-SNE map.

## Supplementary Video 1: Two-photon in vivo calcium imaging of V1 neurons in light-evoked mouse, followed by LS neuron soma extraction

Cal 520-labeled cells in layer 2/3 of mouse V1 were recorded to measure the calcium response to the 5-time repeated light stimulus, under the ‘green’ channel of the two-photon microscope. A light-responding neuron was then targeted to extract the contents of the soma via a glass cuspidal electrode. The light response cells are indicated by yellow arrows, and the target cell is indicated by the red arrow. The frame of calcium imaging was shifted to the left for a better view of the soma extraction. The cells marked with white arrowheads are coordinate references for locating the target cell. Scale bar, 20 µm.

## Supplementary Video 2: Two-photon in vivo whole-cell configuration

Left panel: 5-minute movie of whole-cell patch clamp configuration with the cuspidal electrode under the ‘red’ channel of the two-photon microscope. The shadow of the target cell can be seen. Scale bar, 20 µm. Right panel: the corresponding membrane test monitor from the recording software of Clampex. The cells marked with white arrowheads marked cells are coordinate references for the target cell. Scale bar, 20 µm.

## Supplementary Video 3: In vivo dual recording of light-evoked calcium response and action potential firing

Left panel: the video shows the calcium response to light stimulation of a whole-cell patched neuron (Cal-520 labeled), under the ‘green’ channel of the two-photon microscope. The target neutron is indicted by a yellow arrow and the position of the glass electrode is marked. Scale bar, 20 µm. Right panel: the calcium response and the action potential firing are dynamically represented in the ‘calcium signal’ and ‘electrophysiological signal’ windows, respectively.

## Supplementary Video 4: 3-D morphological reconstruction of a neuron in layer 2/3 of V1

The video is related to Figure 2g. Scale bar, 30 µm.

